# The metabolite receptor SUCNR1 in oxidative stress-induced age-related macular degeneration

**DOI:** 10.1101/2020.02.24.962779

**Authors:** Elja M.M. Louer, Peter M.T. Deen, Anneke I. den Hollander

## Abstract

Age-related macular degeneration (AMD) is the leading cause of vision impairment in elderly people. AMD is a multifactorial disease which is characterised by complex interactions between metabolic and environmental factors as well as multiple genetic susceptibility factors. The exact mechanism of the most prominent environmental factors, age and smoking, in combination with genetic susceptibility factors is little studied. Here, we set out to study the influence of age, smoking induced oxidative stress and the role of succinate receptor 1 (SUCNR1) in AMD development in mice.

*Sucnr1* wild-type (WT), heterozygous (HT) and knock-out (KO) mice were exposed to smoking related oxidative stress by the addition of hydroquinone (HQ), the most abundant oxidant in cigarette smoke, to the drinking water of the mice. Using immunohistochemical staining, accumulation of oxidized LDL (oxLDL) in the mouse retina was assessed at 40 and 48 weeks of age.

At 40 weeks of age, a significant increase in oxLDL in the *Sucnr1* KO mice treated with HQ was observed when compared to the WT and HT mice treated with HQ (p<0.01). However, at 48 weeks, no significant difference was observed between any of the groups. A second experiment analyzing the mice at 40 weeks of age was unable to confirm the observed results of the first experiment.

We identified oxLDL accumulations in *Sucnr1* KO retinas exposed to HQ, but were unable to repeat this finding. Therefore, under the present conditions, the *Sucnr1* KO mouse model is not a suitable model to study AMD development.

## Introduction

Age-related macular degeneration (AMD) is the leading cause of vision impairment in elderly people, and it is estimated that 8.7% of the world population is affected by AMD^1^. AMD is a neurodegenerative disease of the macula and is characterized by the formation of insoluble extracellular lipid aggregates, known as drusen^2,3^. Drusen are found between the retinal pigment epithelium (RPE) layer and the Bruchs’ membrane^4^, and are associated with reduced functionality of the RPE^5^. Based on phenotypic manifestations, advanced AMD can be classified into two distinct forms. Dry AMD is characterized by RPE and photoreceptor (PR) cell death known as geographic atrophy^6^. The second form is called wet AMD due to the neovascularization of the choroid^6^. Understanding the molecular mechanisms underlying wet AMD pathogenesis has resulted in the development of FDA-approved therapies targeting vascular endothelial growth factor to prevent severe visual loss^7^. However, no treatment is currently available yet for the dry form of AMD.

AMD is a multifactorial disease which is characterised by complex interactions between metabolic, functional and environmental factors as well as several genetic susceptibility factors. Moreover, cigarette smoking is indicated as one of the greatest environmental risk factors for the development of both dry and wet AMD^8^. Cigarette smoke contains multiple potential toxic substances, including many pro-oxidants. One of the pro-oxidants found in cigarette smoke is hydroquinone (HQ)^9^, and toxicity of HQ to retinal cells through oxidative, mitochondrial and autophagic pathways has been shown in previous *in vitro* studies^10-12^. Cumulative oxidative damage as a result of an imbalance between the antioxidant defence system and generation of reactive oxygen species (ROS) has long been hypothesized to play a substantial role in RPE impairment and AMD progression^13-16^.

Recently, a potential role for the metabolite sensing receptor succinate receptor 1 (SUCNR1) in AMD development was proposed^17^. It was shown that deficiency of Sucnr1 in mice led to outer retinal lesions and accumulation of oxidative LDL (oxLDL), a lipid-protein complex found in drusen. Moreover, we recently described a single nucleotide polymorphism in the 3’-UTR of the *SUCNR1* gene, which is enriched in the EUGENDA AMD cohort, and reduces the *SUCNR1* expression levels *in vitro*^18^. Understanding the link between the SUCNR1 and the development of AMD may lead to new therapeutic innovations. The objective of this study was to induce smoking-related oxidative stress in mice in absence or presence of the Sucnr1 to document changes in the retinal layers and to observe whether Sucnr1 plays a role in the development of AMD.

## Methods

### Animal studies

*Sucnr1* knock-out mice on a C56BL/6 background were generated as described^19^. *Sucnr1* wild-type (WT), heterozygous (HT) and knock-out (KO) littermates were housed under standardized conditions (12 h dark/12 h light cycle) and fed ad libitum. Animal experiments were approved by the animal ethics board of the Radboud University, Nijmegen (RU DEC 2016-0054) and by the Dutch Central Commission for Animal Experiments (CCD, AVD103002016620).

Eight week-old animals from each genotype group were randomly divided into two groups: untreated controls and HQ-induced oxidative stress. In the oxidative stress groups, continuous stress exposure was induced by addition of 0.8% HQ (Sigma-Aldrich, Burlington, MA, USA) to the animal’s drinking water. At 32, 40 and 48 weeks of age, mice were sacrificed using cervical dislocation and eyes were collected via enucleation. Anterior segments (cornea, iris and lens) were removed to expose the posterior eyecup. Eyes were fixed in 4% PFA at room temperature (RT) followed by graded sucrose embedding (10%, 20% and 30%) (MP Biomedicals, Amsterdam, the Netherlands). Finally, eyecups were enclosed in optimum cutting temperature compound (OCT, Sakura Finetek Holland B.V., Alphen aan den Rijn, the Netherlands) and frozen in liquid nitrogen. Tissue was stored at −80 °C until sectioning.

### Fluorescence immunohistochemistry

Eyecups were cryosectioned at 12 µm thickness using the Microm HM 500 M Cryostat at a chamber temperature of −20°C and a specimen temperature of −17°C. For every eye, six to seven sections were mounted on Superfrost plus slides (ThermoFisher Scientific, Breda, the Netherlands), dried at RT overnight, and stored at −20 °C until staining.

After blocking using 0.5% Blocking Reagent (TSA Fluorescein System kit, PerkinElmer, Cat.No. NEL701A001KT) in Tris/HCl + NaCl, tissue sections were stained with anti-oxLDL (1:400, Sigma-Aldrich, Burlington, MA, USA) overnight at 4°C followed by Alexa 488 (1:300, ThermoFisher Scientific, Breda, the Netherlands) for 2 hours at RT. DAPI Fluoromount-G (SouthernBiotech, Uithoorn, the Netherlands) was used to mount the sections and to stain the nuclei. Fluorescence images were obtained with a Zeiss Imager.M2 fluorescent microscope.

### Image J analysis

Using Image J software^20^ the area of ox-LDL was measured. The total area containing accumulation of ox-LDL for each eye was averaged as the representative value for that eye. The average ox-LDL accumulation of both eyes of a mouse were taken as one measurement. Examiners were masked for the genotype group and treatment regime of the mice during microscopy and ImageJ analysis.

### Statistical analyses

Statistical analyses were performed using Graphpad Prism 8 software. Three-way ANOVA or two-way ANOVA were used, as indicated in figure legends. P-values < 0.05 (*), < 0.01 (**) and < 0.001 (***) were considered to be statistically significant. For the three-way ANOVA, missing values (4 out of 96) were supplemented with the mean of the group for statistical analysis purposes.

### RPE RNA isolation

For RNA-sequencing analysis, 40 week-old control and treated WT and *Sucnr1* KO mice were randomly selected for RNA-sequencing analysis (n=4 per group). Mice were sacrificed at 2 am using cervical dislocation and eyes were collected via enucleation. Anterior segments were removed and the retina was carefully peeled off. The eyecup containing the RPE, sclera and choroid were stored in RNAlater solution (Thermofisher, Amsterdam, The Netherlands) at −20°C.

For RNA isolation, the RPE was removed gently from the choroidal layer. Afterwards, the RPE was dissolved in RLT Buffer from RNeasy Micro Kit (Qiagen, Germantown, MD, USA). RNA was extracted using the RNeasy Micro Kit according to the manufacturer’s protocol using the on column DNAse treatment. RNA concentration and quality were measured using a Nanodrop spectrophotometer (NanoDrop 2000C, Thermofisher, Amsterdam, The Netherlands).

### RNA-sequencing

RNA samples of one eye per mouse were sent to ServiceXS (Genomescan, Leiden, The Netherlands) for sequencing. The RNA concentration and quality were assessed using the Advanced Analytical Fragment Analyzer (Agilent Technologies, CA, USA, table 1) prior to library construction. Library prep was performed using NEBNext Ultra Directional RNA Library Prep Kit from Illumina (New England Biolabs, Leiden, The Netherlands). 1.1 nM of cDNA was used for input samples in Illumina Novasec 6000 sequencer (Illumina, San Diego, USA) for 1 × 150 bp paired-end sequencing, sequencing an average of 20 million reads per sample.

**Table 1.**
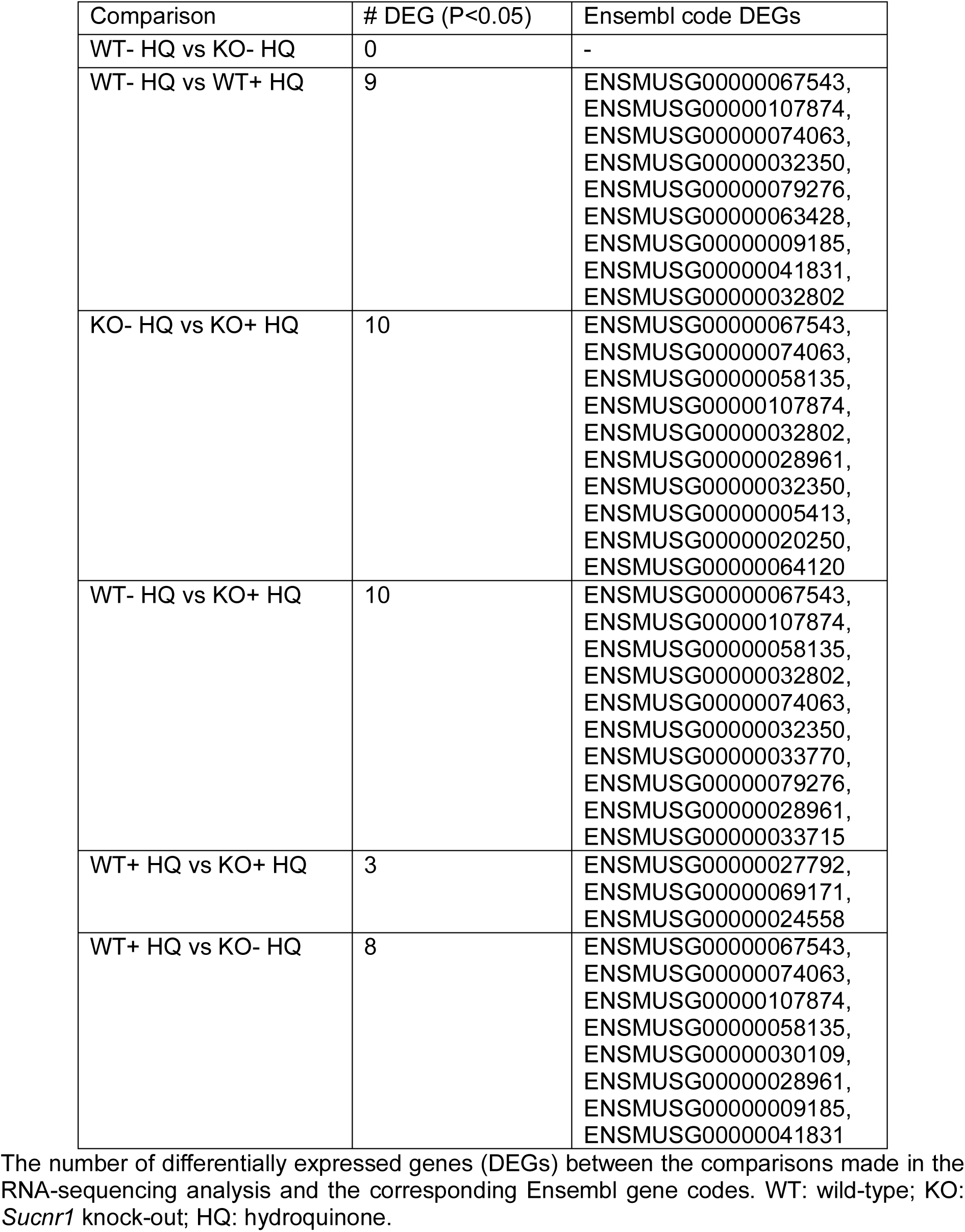
Number of identified differentially expressed genes.

### RNA-sequencing data analyses

The mm10 reference assembly was first indexed by STAR mapper^21^ with gene annotation model from ENSEMBL database. RNA-seq reads were aligned to the mm10 genome using the two-pass mode of STAR, and the gene-based read counts were quantified at the same time. The DESeq2 package^22^ was used to detect differentially expressed genes by performing pair-wise comparison among the four described time points. Only genes with a Benjamini-Hochberg-adjusted p-value < 0.05 and fold change > 1.5 were considered significantly differentially expressed. Cufflinks^23^ was employed to estimate gene-level abundance in FPKM values for ENSEMBL genes.

### Data availability

The RNA-seq data used in this paper has been deposited in the Gene Expression Omnibus database^24^ with the accession number GSE139996.

## Results

To study the combined effect of absence or reduced *Sucnr1* expression and HQ-induced oxidative stress on AMD development, *Sucnr1* WT, HT and KO mice were used. Mice from each genotype group were either exposed to HQ from the age of eight weeks or left untreated, resulting in six experimental groups. Each experimental group contained eight mice for each time point of tissue isolation. At 32, 40 and 48 weeks of age, eyes were quantitatively analyzed for drusen formation using anti-oxLDL staining, and for each mouse the area of oxLDL staining of both eyes was averaged.

Representative images of anti-oxLDL staining are shown in figure 1. At 32 weeks of age, no specific anti-oxLDL staining was observed (data not shown). At 40 weeks of age, an increase in oxLDL was observed in the KO animals treated with HQ. After quantification, a significant increase in oxLDL staining was observed between the HQ-treated WT compared to HQ-treated KO animals (p=0.0061), between the HQ-treated HT compared to HQ-treated KO animals (p=0.009) and between untreated KO compared to HQ-treated KO animals (p=0.0057; figure 2A). Moreover, the HT animals treated with HQ seem to show an intermediate phenotype compared to the HQ-treated WT and KO animals.

**Figure 1.**
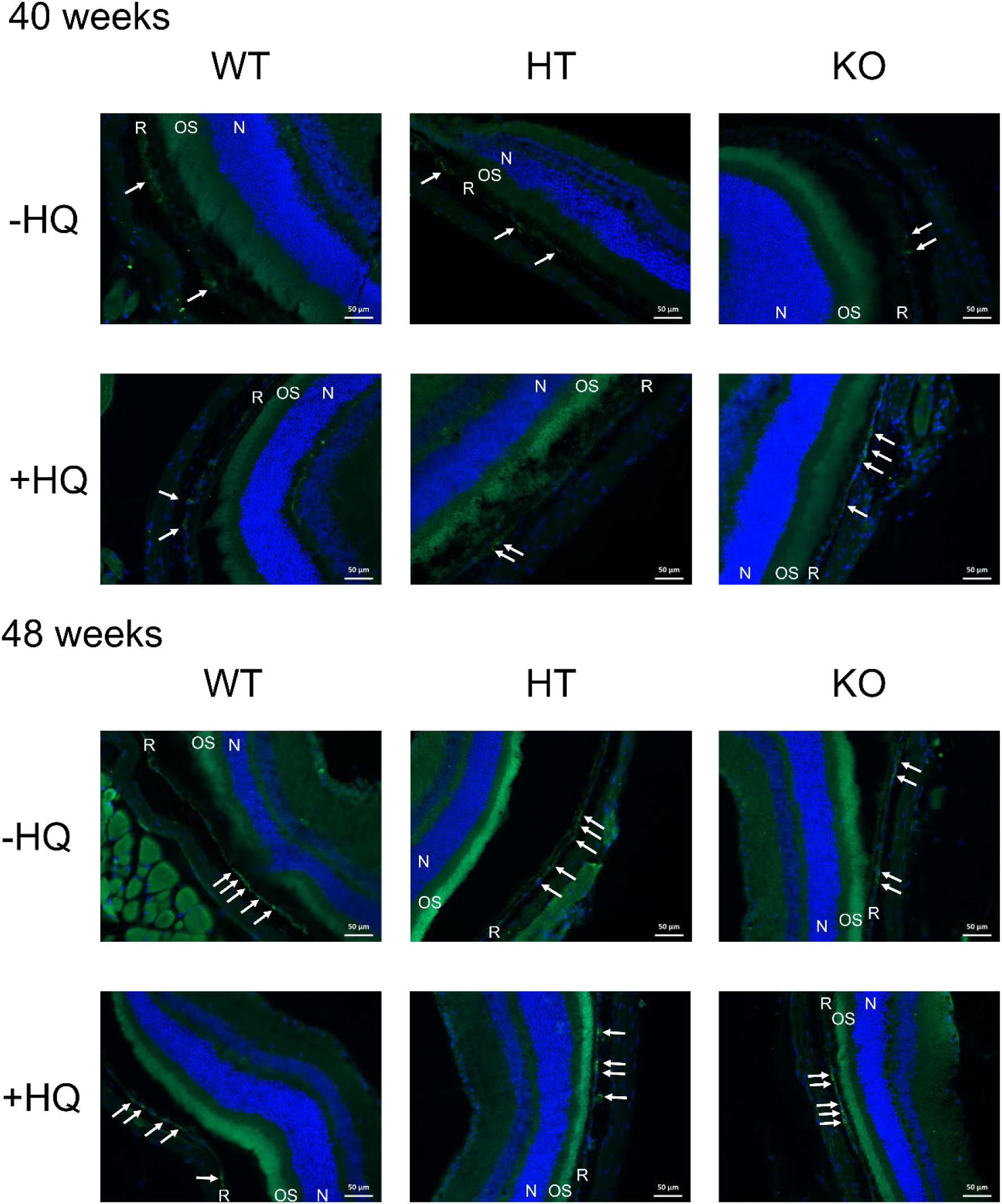
Anti-oxLDL staining of retinal sections from 40- and 48-week-old mice. Representative pictures of anti-oxLDL staining (in green) at 40 and 48 weeks of age for the 6 experimental groups are shown with DAPI stained in blue. Specific staining in the RPE is indicated with the white arrows. Scale bars: 50 µm. N: outer nuclear layer; OS: outer segments; R: RPE; HQ: hydroquinone-treated; WT: wild-type; HT: *Sucnr1* heterozygote; KO: *Sucnr1* knock-out.

**Figure 2.**
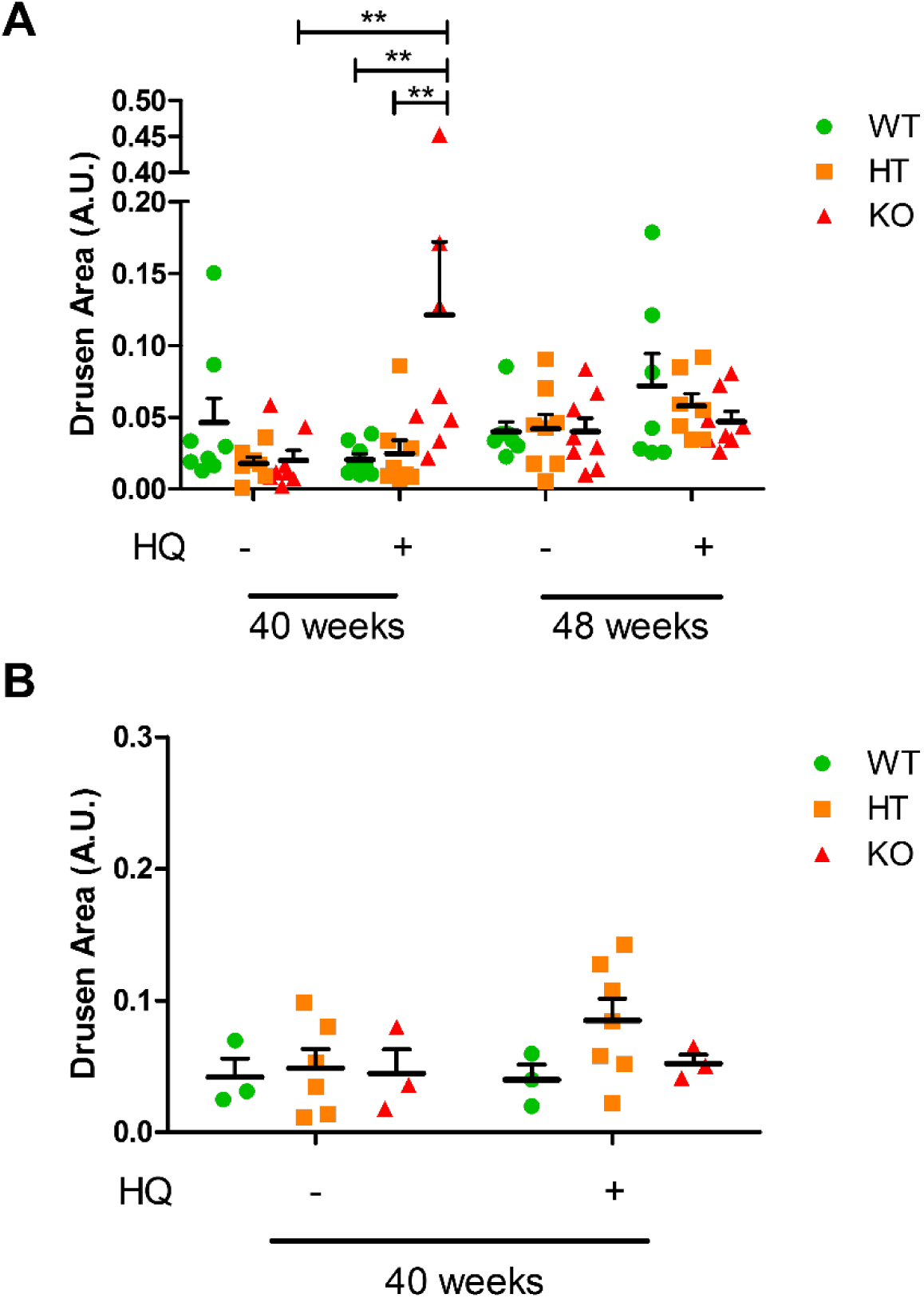
Quantification of anti-oxLDL staining. Anti-oxLDL staining was quantified using ImageJ software. Average amount of staining per mouse is plotted (n=7-8). A) Anti-oxLDL staining measured for first animal experiment at 40 and 48 weeks of age. Data analyzed using three-way ANOVA. **, p<0.01. B) Anti-oxLDL staining measured in an independent second animal experiment at 40 weeks of age. Data analyzed using two-way ANOVA. HQ: hydroquinone-treated; WT: wild-type; HT: *Sucnr1* heterozygote; KO: *Sucnr1* knock-out.

At 48 weeks of age the anti-oxLDL staining showed no significant difference between the groups. However, a trend for increased oxLDL staining was observed when comparing all treated groups to their untreated control groups.

To further study and verify the effect observed at 40 weeks of age, the same genotype groups were investigated in an independent second experiment (n=7 per experimental group). RNA was isolated from WT and KO eyes from both the treated and untreated groups, which was subjected to RNA sequencing (n=4 per experimental group). When comparing all groups to each other, very little significant differences in gene expression were observed (table 1). The remaining animals were used for anti-oxLDL staining (n=3-7). After quantification of the anti-oxLDL staining, no difference was observed between the six experimental groups (figure 2B).

## Discussion

This study sought to identify and elucidate the role of SUCNR1 deficiency in oxidative stress-induced development of (dry) AMD. Previously, Favret et al. showed that 43 week-old *Sucnr1* KO mice developed thickening of the Bruch’s membrane compared to aged *Sucnr1* WT mice^17^. Moreover, they showed oxLDL accumulations in *Sucnr1* KO mice of the same age exposed to a high fat diet from the eighth week of life. Due to the multifactorial aspect of AMD development, we set out to study if smoking-induced oxidative stress could induce similar effects on oxLDL deposit formation. Furthermore, we intended to study affected pathways underlying SUCNR1 deficiency related to AMD development.

The different results between our study and that of Favret et al. are striking. In our first experiment, we used eight mice per experimental group to quantity oxLDL accumulation where Favret et al. used three to four mice. The mice showing oxLDL accumulations in the study of Favret et al. were 43 weeks old KO animals and were exposed to a high fat diet. We first detected oxLDL accumulations in 40 week-old KO mice exposed to HQ. In contrast to Favret et al., our anti-oxLDL staining was quantitatively evaluated using imageJ software, while Favret et al. did not perform a quantitative analysis of oxLDL staining. Our data revealed a significant increase in oxLDL in the *Sucnr1* KO mice treated with HQ which is in line with the results of Favret et al. in *Sucnr1* KO mice exposed to a high fat diet. However, we were not able to repeat our first finding in an independent second experiment where we used three to seven animals per experimental group. Moreover, sequencing of RPE RNA from WT and *Sucnr1* KO animals both untreated and treated with HQ did not reveal many differentially expressed genes.

The fact that we could not repeat our first findings may be explained by the multifactorial nature of AMD. Oxidative stress has long been considered as a major contributor to the pathophysiology of AMD^8^. However, the pathophysiology of AMD is complex, and multiple processes are considered to contribute to disease progression. Within our experiments, experimental groups only varied in the presence of oxidative stress induced by HQ treatment and the presence of Sucnr1. Oxidative stress induced by other factors, such as inflammation, outer-segment phagocytosis, aging, and photo-oxidative stress from processing light for vision was equal in all experimental groups^8^. Therefore, it is likely that Sucnr1 deficiency and HQ treatment are only minor contributors to the overall disease process whereas the high fat diet used by Favret et al. has bigger implications.

A genome-wide association study (GWAS) using over 16,000 AMD patients and 17,000 controls have linked AMD to several genes involved in lipid metabolism and the complement system^25^. Favret et al. described several genetic variants in the *SUCNR1* gene to be associated with dry AMD^17^ and in a recent study we also identified a genetic variant in the *SUCNR1* gene to be associated with AMD^18^. Furthermore, we showed a possible mechanism of *SUCNR1* mRNA downregulation as a result of this variant. However, the GWAS study in 16,000 AMD patients and 17,000 control individuals did not find any significant associations for genetic variants in the *SUCNR1* gene^25^. This further strengthens the suggestion that the contribution of SUCNR1 deficiency to the development of AMD is only small, or that other contributing factors were not taken into account during the analyses.

To conclude, we identified oxLDL accumulations in *Sucnr1* KO retinas exposed to HQ, but were unable to repeat this finding in an independent experiment. Moreover, after critically evaluating the data from Favret et al. we suggest that the role of SUCNR1 deficiency in AMD development is only small. Therefore, under the present conditions the *Sucnr1* KO mouse model is not a suitable model to study AMD development.

## Acknowledgements

Anneke I. den Hollander and Peter M.T. Deen are co-corresponding authors for this article. We thank Valerie Villacorta Monge, Maud Wekking and Juul Verbakel for their technical assistance. This study was supported by a junior researcher grant from the Radboud Institute for Molecular Life Sciences and Radboudumc to Peter M.T. Deen and Anneke I. den Hollander (RIMLS014-056).

## References

1 Wong, W. L., Su, X., Li, X., Cheung, C. M., Klein, R., Cheng, C. Y. & Wong, T. Y. Global prevalence of age-related macular degeneration and disease burden projection for 2020 and 2040: a systematic review and meta-analysis. Lancet Glob Health 2, e106–116, doi:10.1016/S2214-109X(13)70145-1 (2014).

2 Wang, L., Clark, M. E., Crossman, D. K., Kojima, K., Messinger, J. D., Mobley, J. A. & Curcio, C. A. Abundant lipid and protein components of drusen. PLoS One 5, e10329, doi:10.1371/journal.pone.0010329 (2010).

3 Bergen, A. A., Arya, S., Koster, C., Pilgrim, M. G., Wiatrek-Moumoulidis, D., van der Spek, P. J., Hauck, S. M., Boon, C. J. F., Emri, E., Stewart, A. J. & Lengyel, I. On the origin of proteins in human drusen: The meet, greet and stick hypothesis. Prog Retin Eye Res 70, 55–84, doi:10.1016/j.preteyeres.2018.12.003 (2019).

4 Russell, S. R., Mullins, R. F., Schneider, B. L. & Hageman, G. S. Location, substructure, and composition of basal laminar drusen compared with drusen associated with aging and age-related macular degeneration. Am J Ophthalmol 129, 205–214, doi:10.1016/s0002-9394(99)00345-1 (2000).

5 Hageman, G. S., Luthert, P. J., Victor Chong, N. H., Johnson, L. V., Anderson, D. H. & Mullins, R. F. An integrated hypothesis that considers drusen as biomarkers of immune-mediated processes at the RPE-Bruch’s membrane interface in aging and age-related macular degeneration. Prog Retin Eye Res 20, 705–732 (2001).

6 Jager, R. D., Mieler, W. F. & Miller, J. W. Age-related macular degeneration. N Engl J Med 358, 2606–2617, doi:10.1056/NEJMra0801537 (2008).

7 Rosenfeld, P. J., Brown, D. M., Heier, J. S., Boyer, D. S., Kaiser, P. K., Chung, C. Y., Kim, R. Y. & Group, M. S. Ranibizumab for neovascular age-related macular degeneration. N Engl J Med 355, 1419–1431, doi:10.1056/NEJMoa054481 (2006).

8 Evans, J. R. Risk Factors for Age-related Macular Degeneration. Prog. Retin. Eye Res. 20, 227–253, doi: https://doi.org/10.1016/S1350-9462(00)00023-9 (2001).

9 Dagnon, S., Stoilova, A., Ivanov, I. & Nikolova, S. The Effect of Cigarette Design on the Content of Phenols in Mainstream Tobacco Smoke. Vol. 24 (2010).

10 Sharma, A., Patil, J. A., Gramajo, A. L., Seigel, G. M., Kuppermann, B. D. & Kenney, C. M. Effects of hydroquinone on retinal and vascular cells in vitro. Indian J. Ophthalmol. 60, 189–193, doi:10.4103/0301-4738.95869 (2012).

11 Ramírez, C., Pham, K., Franco, M. F. E., Chwa, M., Limb, A., Kuppermann, B. D. & Kenney, M. C. Hydroquinone induces oxidative and mitochondrial damage to human retinal Müller cells (MIO-M1). Neurotoxicology 39, 102–108, doi: https://doi.org/10.1016/j.neuro.2013.08.004 (2013).

12 Espinosa-Heidmann, D. G., Suner, I. J., Catanuto, P., Hernandez, E. P., Marin-Castano, M. E. & Cousins, S. W. Cigarette smoke-related oxidants and the development of sub-RPE deposits in an experimental animal model of dry AMD. Invest Ophthalmol Vis Sci 47, 729–737, doi:10.1167/iovs.05-0719 (2006).

13 Fanjul-Moles, M. L. & Lopez-Riquelme, G. O. Relationship between Oxidative Stress, Circadian Rhythms, and AMD. Oxid Med Cell Longev 2016, 7420637, doi:10.1155/2016/7420637 (2016).

14 Bowes Rickman, C., Farsiu, S., Toth, C. A. & Klingeborn, M. Dry age-related macular degeneration: mechanisms, therapeutic targets, and imaging. Invest. Ophthalmol. Vis. Sci. 54, ORSF68–ORSF80, doi:10.1167/iovs.13-12757 (2013).

15 Ambati, J. & Fowler, B. J. Mechanisms of age-related macular degeneration. Neuron 75, 26–39, doi:10.1016/j.neuron.2012.06.018 (2012).

16 Datta, S., Cano, M., Ebrahimi, K., Wang, L. & Handa, J. T. The impact of oxidative stress and inflammation on RPE degeneration in non-neovascular AMD. Prog Retin Eye Res 60, 201–218, doi:10.1016/j.preteyeres.2017.03.002 (2017).

17 Favret, S., Binet, F., Lapalme, E., Leboeuf, D., Carbadillo, J., Rubic, T., Picard, E., Mawambo, G., Tetreault, N., Joyal, J. S., Chemtob, S., Sennlaub, F., Sangiovanni, J. P., Guimond, M. & Sapieha, P. Deficiency in the metabolite receptor SUCNR1 (GPR91) leads to outer retinal lesions. Aging (Albany NY) 5, 427–444, doi:10.18632/aging.100563 (2013).

18 Louer, E. M. M., Lores-Motta, L., Ion, A. M., Den Hollander, A. I. & Deen, P. M. T. Single nucleotide polymorphism rs13079080 is associated with differential regulation of the succinate receptor 1 (SUCNR1) gene by miRNA-4470. RNA Biol 1-8, doi:10.1080/15476286.2019.1643100 (2019).

19 van Diepen, J. A., Robben, J. H., Hooiveld, G. J., Carmone, C., Alsady, M., Boutens, L., Bekkenkamp-Grovenstein, M., Hijmans, A., Engelke, U. F. H., Wevers, R. A., Netea, M. G., Tack, C. J., Stienstra, R. & Deen, P. M. T. SUCNR1-mediated chemotaxis of macrophages aggravates obesity-induced inflammation and diabetes. Diabetologia 60, 1304–1313, doi:10.1007/s00125-017-4261-z (2017).

20 Schneider, C. A., Rasband, W. S. & Eliceiri, K. W. NIH Image to ImageJ: 25 years of image analysis. Nat Methods 9, 671–675 (2012).

21 Dobin, A., Davis, C. A., Schlesinger, F., Drenkow, J., Zaleski, C., Jha, S., Batut, P., Chaisson, M. & Gingeras, T. R. STAR: ultrafast universal RNA-seq aligner. Bioinformatics 29, 15–21, doi:10.1093/bioinformatics/bts635 (2013).

22 Love, M. I., Huber, W. & Anders, S. Moderated estimation of fold change and dispersion for RNA-seq data with DESeq2. Genome Biol 15, 550, doi:10.1186/s13059-014-0550-8 (2014).

23 Trapnell, C., Williams, B. A., Pertea, G., Mortazavi, A., Kwan, G., van Baren, M. J., Salzberg, S. L., Wold, B. J. & Pachter, L. Transcript assembly and quantification by RNA-Seq reveals unannotated transcripts and isoform switching during cell differentiation. Nat Biotechnol 28, 511–515, doi:10.1038/nbt.1621 (2010).

24 Edgar, R., Domrachev, M. & Lash, A. E. Gene Expression Omnibus: NCBI gene expression and hybridization array data repository. Nucleic Acids Res 30, 207–210, doi:10.1093/nar/30.1.207 (2002).

25 Fritsche, L. G., Igl, W., Bailey, J. N., Grassmann, F., Sengupta, S., Bragg-Gresham, J. L., Burdon, K. P., Hebbring, S. J., Wen, C., Gorski, M., Kim, I. K., Cho, D., Zack, D., Souied, E., Scholl, H. P., Bala, E., Lee, K. E., Hunter, D. J., Sardell, R. J., Mitchell, P., Merriam, J. E., Cipriani, V., Hoffman, J. D., Schick, T., Lechanteur, Y. T., Guymer, R. H., Johnson, M. P., Jiang, Y., Stanton, C. M., Buitendijk, G. H., Zhan, X., Kwong, A. M., Boleda, A., Brooks, M., Gieser, L., Ratnapriya, R., Branham, K. E., Foerster, J. R., Heckenlively, J. R., Othman, M. I., Vote, B. J., Liang, H. H., Souzeau, E., McAllister, I. L., Isaacs, T., Hall, J., Lake, S., Mackey, D. A., Constable, I. J., Craig, J. E., Kitchner, T. E., Yang, Z., Su, Z., Luo, H., Chen, D., Ouyang, H., Flagg, K., Lin, D., Mao, G., Ferreyra, H., Stark, K., von Strachwitz, C. N., Wolf, A., Brandl, C., Rudolph, G., Olden, M., Morrison, M. A., Morgan, D. J., Schu, M., Ahn, J., Silvestri, G., Tsironi, E. E., Park, K. H., Farrer, L. A., Orlin, A., Brucker, A., Li, M., Curcio, C. A., Mohand-Said, S., Sahel, J. A., Audo, I., Benchaboune, M., Cree, A. J., Rennie, C. A., Goverdhan, S. V., Grunin, M., Hagbi-Levi, S., Campochiaro, P., Katsanis, N., Holz, F. G., Blond, F., Blanche, H., Deleuze, J. F., Igo, R. P., Jr., Truitt, B., Peachey, N. S., Meuer, S. M., Myers, C. E., Moore, E. L., Klein, R., Hauser, M. A., Postel, E. A., Courtenay, M. D., Schwartz, S. G., Kovach, J. L., Scott, W. K., Liew, G., Tan, A. G., Gopinath, B., Merriam, J. C., Smith, R. T., Khan, J. C., Shahid, H., Moore, A. T., McGrath, J. A., Laux, R., Brantley, M. A., Jr., Agarwal, A., Ersoy, L., Caramoy, A., Langmann, T., Saksens, N. T., de Jong, E. K., Hoyng, C. B., Cain, M. S., Richardson, A. J., Martin, T. M., Blangero, J., Weeks, D. E., Dhillon, B., van Duijn, C. M., Doheny, K. F., Romm, J., Klaver, C. C., Hayward, C., Gorin, M. B., Klein, M. L., Baird, P. N., den Hollander, A. I., Fauser, S., Yates, J. R., Allikmets, R., Wang, J. J., Schaumberg, D. A., Klein, B. E., Hagstrom, S. A., Chowers, I., Lotery, A. J., Leveillard, T., Zhang, K., Brilliant, M. H., Hewitt, A. W., Swaroop, A., Chew, E. Y., Pericak-Vance, M. A., DeAngelis, M., Stambolian, D., Haines, J. L., Iyengar, S. K., Weber, B. H., Abecasis, G. R. & Heid, I. M. A large genome-wide association study of age-related macular degeneration highlights contributions of rare and common variants. Nat Genet 48, 134–143, doi:10.1038/ng.3448 (2016).

